# Geostatistical visualization of ecological interactions in tumors

**DOI:** 10.1101/729368

**Authors:** Hunter Boyce, Parag Mallick

## Abstract

Recent advances in our understanding sof cancer progression have highlighted the important role played by proteomic heterogeneity and the tumor microenvironment. Single-cell measurement technologies have enabled deep investigation of tumor heterogeneity. However, tools to visualize and interpret single-cell data have lagged behind experimental methods; currently, dimensionality reduction and clustering techniques, such as *t*-sne and SPADE, are the most prevalent visualization techniques. However, such techniques do not enable the visualization of the super-cellular structures that arise either via microenvironmental forces (e.g. hypoxia) or through cell-cell interactions. They additionally do not allow the visualization of ecological niches that are critical to understanding tumor behavior. The advent of novel experimental hyperplex immunostaining platforms, capable of measuring the *in situ* protein expression of dozens of proteins simultaneously, necessitates novel computational tools to quantify and visualize spatial patterns in the tumor microenvironment. Here, we introduce an approach to visualize tumor heterogeneity that integrates multiple geostatistics to capture both global and local spatial patterns. We assess the utility of this approach using an agent-based model of tumor growth under four ecological contexts; predation, mutualism, commensalism, and parasitism. Spatial patterns are visualized in real time as the models progress. This application introduces both an ecological framework for characterizing cellular interactions in cancer and novel way of quantifying and visualizing spatial patterns in cancer.

## 1. Introduction

Recent reports have demonstrated that intratumoral-heterogeneity, including genetic mutation, expression noise, and microenvironmental variation can have a significant impact tumor progression and invasion.^1–3^ Single-cell RNA sequencing^4^ and technologies such as mass cytometry,^5^ are a promising solution to profiling heterogeneity within the tumor by quantifying the RNA content within individual cells or by surveying portions of the cellular proteome. Analysis of this data type attempts to visualize cellular phenotypes and their lineages through dimensionality reduction techniques like t-stochastic neighbor embedding (t-SNE),^6^ and spanning-tree progression analysis of density-normalized events (SPADE).^7^ Despite being extremely powerful tools to understand heterogeneity, they do not provide any information about cell’s context or about what that context may imply about the ecological and evolutionary forces driving tumor behavior.

Recent platforms^8–10^ have emerged that can capture the expression of dozens of proteins and RNA *in situ* in a high-throughput fashion. These tools make it possible to go beyond measuring molecular heterogeneity to understanding spatial relationships amongst diverse cell types and molecularly defined subtypes. With the ability to survey the spatial context of cells within tissues comes computational challenges to understand how to quantify spatial interactions and how to appropriately visualize the intercellular and higher-order spatial relationships. For example, identifying regions where expression of a protein of interest is concentrated or to identify sudden spatial shifts in protein expression.

Visualizing spatial phenomena in tumors has predominantly been limited to protein expression, cell type, or a single geostatistical measure.^11–15^ While these measures have been shown to have modest clinical utility, they do not readily provide insight into underlying biological processes. We hypothesize that jointly monitoring six geostatistics will enable visualization of spatial processes at both the global and local scale. To test this hypothesis, we use agent-based models to grow virtual tumors exhibiting four different ecological processes: predation, mutualism, commensalism, and parasitism. Using our new visualization approach to simultaneously examine protein expression, and global and local spatial phenomena, we demonstrate how a multi-parametric approach enables a more complete view of the spatial ecology of the tumor.

## 2. Background

### Tumor ecology

The interactions between the diverse types of cells within a tumor (epithelial, stromal, immune) and their environment have multiple analogues in community ecology and can be described by four classes of interactions which have all been reported to exist within tumors:^16–18^ predation, mutualism, commensalism, and parasitism. Predatory interactions consist of those where one organism targets and kills another organism. In tumors, this behavior is most prominently noted between certain immune cells that target and kill cancer cells.^19^ Research suggests that the degree of co-localization between these two cells types can influence the progression of the tumor.^12,15^ Mutualistic relationships are those in which two species both benefit by interacting with one another. This type of cooperation has been observed among tumor subclones that harbored mutations which mutualistically created an environment that favored tumor growth.^20^ Commensalism is described as relationships where one species benefits from interaction with a second species that is neither benefited nor harmed. Like mutualism, commensal interactions have been observed between tumor subclones wherein one subclone benefits from consuming a substrate that the other produces.^21^ Lastly, parasitic interactions in which one species benefits from harming, but not killing, another species. In cancer, this is observed when tumor cells steer high energy substrates away from nearby healthy tissues in the reverse Warburg effect.^22^

## 3. Methods

### 3.1. Agent-based Modeling

A spatial, agent-based model was used to simulate a basic tumor growing. The model considers two-dimensional space as a hexagonal grid where each position has six direct neighbors with which it can interact (Figure 1a). The model keeps track of the number of agents and the width and height of the canvas. The canvas is non-toroidal, such that the top and bottom, and left and right, edges of the canvas do not wrap to each other. The model employs a random activation schedule wherein agents are activated in a random order at each step of the model.

**Fig. 1.**
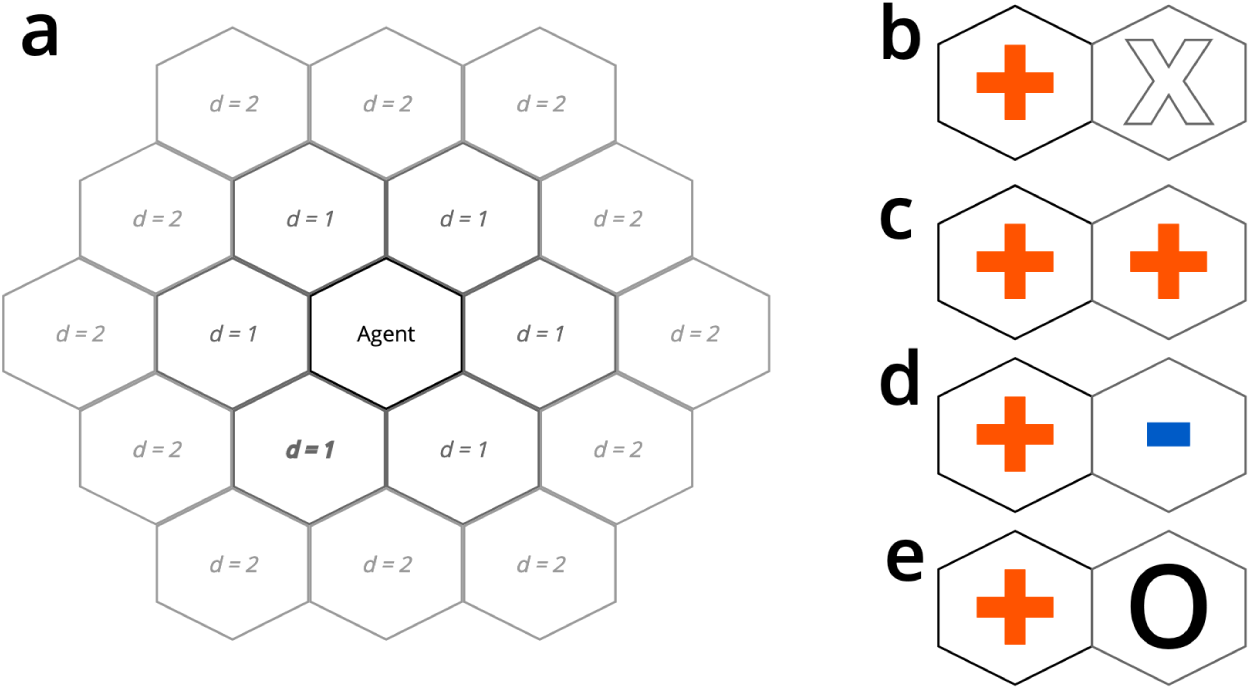
Agent-based model neighbors and interactions. (a) Any agents within a hexagonal distance *d* = 2 are considered neighbors of the central agent. The change in oncoprotein expression upon ecological interaction in (b) predation, (c) mutualism, (d) parasitism, and (e) commensal models.

Each agent in the model represents a cell in the tumor microenvironment. The cell can be a cancer cell, a cancer stem cell, or an immune cell. Each cell has a unique id, a hexagonal coordinate position in the model, an instantaneous death probability, and a division probability. Cancer cells have an oncoprotein parameter which represents the measurement of Ki67, a biomarker of proliferative capacity. Cancer cells also have a counter for the number of divisions remaining before cell death. Cancer stem cells act like cancer cells, except that they have replicative immortality and therefore have an infinite number of divisions remaining. Immune cells have a probability to target a neighboring cancer cell and a probability to kill the cancer cell it is targeting.

The model is initialized with an initial number of cancer cells and a width and height for the canvas. Experimental pathology samples show strong spatial autocorrelation of protein abundance. This necessitates a sampling strategy that considers neighbors for the oncoprotein measurement of the cells. To achieve this we utilize a Markov Chain Monte Carlo (MCMC) sampling strategy:

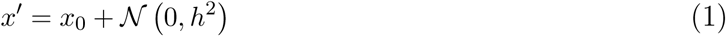

Where *x*_0_ is the mean oncoprotein of your neighbors and *h* is the walk rate for the MCMC. We use the Metropolis-Hastings algorithm to step through the chain where we calculate an acceptance probability:

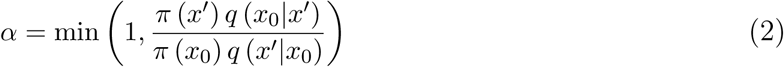

Where *q* is the proposal distribution and *π* is the full joint density. We then sample a uniform random variable *u* and if *u < α*, we accept the proposal *x*′, otherwise we reject and sample the MCMC again. We select a walk rate *h* for the MCMC such that the proposal acceptance rate is around 90% giving an autocorrelated sample. This sampling strategy is used in the initialization of the model and when new agents are added to the model. After oncoprotein initialization for each agent, its fitness is modified using a piecewise generalized logistic function:

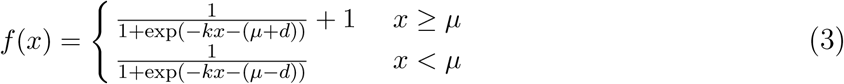

Where *x* is the oncoprotein of the cell, *k* is the growth rate of the logistic curve, *µ* is the mean of the target distribution from the MCMC sampling, and *d* is how far away from *µ* that growth occurs. The resulting *f* (*x*) is used as a coefficient to the division and death probabilities for the agent.

### 3.2. Modeling Ecological Interactions

To model predatory interactions (Figure 1b) in the tumor microenvironment, we initialize the tumor model with cancer cells and cancer stem cells in the center of the model canvas and initialize a perimeter of immune cells encircling the tumor. Immune cells are inactive until the tumor reaches a cancer cell count threshold, upon which the immune cells are activated. While the tumor continues to grow, immune cells begin to move with preference toward the weighted center of the cancer cells. When an immune cell neighbors a cancer cell it will target and attach to the cancer with its initialized target probability. If an immune cell is already targeting a cancer cell, it will kill that cell with its initialized kill probability. Immune cells can only target one cancer cell at a time and targeting and killing occur at different steps of the model.

To model mutualistic interactions (Figure 1c) in the tumor microenvironment, we initialize two separate tumors on the canvas, each with cancer cells and cancer stem cells, and the tumors are allowed to grow. If cancer cells from the first tumor neighbor cancer cells from the second tumor, a mutualistic interaction occur and both cells increase in oncoprotein expression and gain a fitness benefit. This allows greater replicative capacity. This increase in proliferative ability is not compound and is only allowed to happen once.

Modeling commensal interactions (Figure 1d) in the tumor microenvironment is initialized in the same manner as the mutualism model. The difference being, when cancer cells from the first tumor neighbor cancer cells from the second, the first gains a fitness advantage while the other is unchanged. Like the mutualism model, the fitness advantage is not compound.

The parasitic model (Figure 1e) is setup in the same way as the mutualism and commensalism models. However, when cancer cells from the first tumor neighbor cancer cells from the second, the first gains a fitness advantage while the other has a decrease in proliferative capability. Similar to the previous two models, the change in proliferative capacity is not compound.

### 3.3. Geostatistical Methods

For all equations in this section, *x*_*i*_ is the oncoprotein expression of cell *i* and *w*_*ij*_ = 1 if cell *i* and cell *j* are neighbors in the model. Otherwise *w*_*ij*_ = 0. Cells are considered neighbors if they have a Manhattan distance of two or less (Figure 1a).

#### 3.3.1. Global Statistics

Global statistics are geostatistics describing the entire model at the current step. We calculate three global statistics at each step that describe how oncoprotein is clustered and the spatial heterogeneity of oncoprotein in the model. To quantify general clustering of oncoprotein at the global level we use the Getis-Ord general *G* statistic:^23^

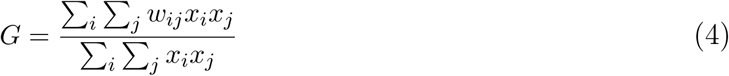

The *G* statistic is bounded between 0, indicating clustering of low values, and 1 indicating clustering of high values. To quantify spatial heterogeneity we utilize two statistics. First, Moran’s *I* statistic^24^ :

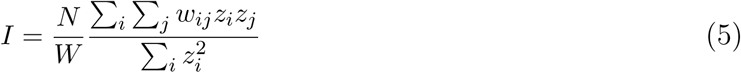

Where *N* is the number of agents in the model, *W* = Σ _*i*_ Σ _*j*_ *w*_*ij*_, and 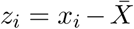 The value of *I* is between *-*1 for perfectly dispersed (low autocorrelation) data and +1 for perfectly clustered (high autocorrelation) data. The second statistic that we use to quantify spatial heterogeneity is Geary’s C^25^ :

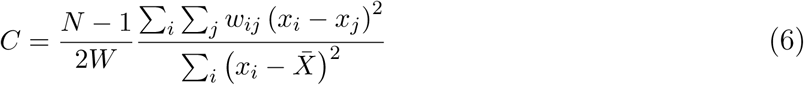

Where *N* is the number of agents in the model and *W* = Σ_*i*_ Σ_*j*_ *w*_*ij*_. Geary’s *C* is inversely related to Moran’s *I* and can take on values between 0 with perfect clustering (high autocorrelation) and an undefined upper bound for increased dispersion (low autocorrelation.

#### 3.3.2. Local Statistics

Local geostatistics describe spatial behavior at the agent level. Each of the global geostatistics has a local counterpart. However, instead of having a single statistic for the entire model, each agent in the model takes on a value for these statistics. To quantify the degree of spatial clustering we utilize the local Getis-Ord *G*_*i*_ statistic^23,26^ :

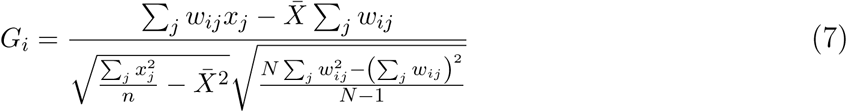

Large positive values of *G*_*i*_ correspond to a hotspot of expression at location *i* and low negative values correspond to a coldspot. To quantify local spatial heterogeneity we again use two different statistics. First, Anselin local Moran’s *I*_*i*_^24,27^ :

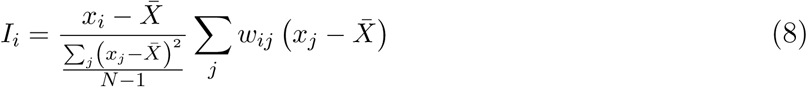

A positive value for *I*_*i*_ indicates that neighbors of agent *i* are similar to itself (low heterogeneity). A negative value indicates dissimilar neighbors (high heterogeneity). The second statistic for spatial heterogeneity is the local Geary’s *C*_*i*_ statistic^25,27^ :

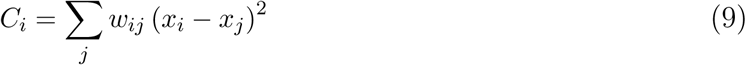

A value close to zero for *C*_*i*_ indicates homogeneous neighbors and large positive values indicate heterogeneous neighbors.

## 4. Results and Discussion

An ideal approach for measuring and visualizing spatial heterogeneity in tumors should have the following qualities: 1) sensitive to true variation in a growing tumor over time ; 2) insensitive minor variations in a broadly unchanging tumor; 3) does not present multiple dimensions of redundant information; and 4) differentiates among distinct ecological interactions. Below we examine the performance of our approach to measure and visualize tumor ecology and heterogeneity across a series of virtual tumors exhibiting a range of ecological patterns.

### 4.1. Predatory behavior results in continually varying geostatistics

Visualizing the global behavior of the predation model (Figure 2a-d) shows a global response that is very sensitive to the state of the model. In the initial stages of the model, the tumor is allowed to grow unimpeded with no interaction with immune cells. This is behavior is reflected in global geostatistics that remain stable during this initial period. As the tumor grows, ΔN steadily grows relative to the size of the tumor until predatory interspecific interactions occur and growth drops precipitously at step 160 of the model (Figure 2a). This is met with an increase spatial heterogeneity as immune cells are interfacing with cancer cells (Figure 2b,f). As the immune cells successfully kill and target cancer cells through step 260, we observe a decrease in spatial heterogeneity and an increase in spatial clustering as marked by a decrease in *C* and an increase in *G* and *I* (Figure 2b-d). This can be explained by a spatially constrained cancer cell population with high autocorrelation. When the immune cells target the remaining cancer cells after step 260, we observe another increase in spatial heterogeneity.

**Fig. 2.**
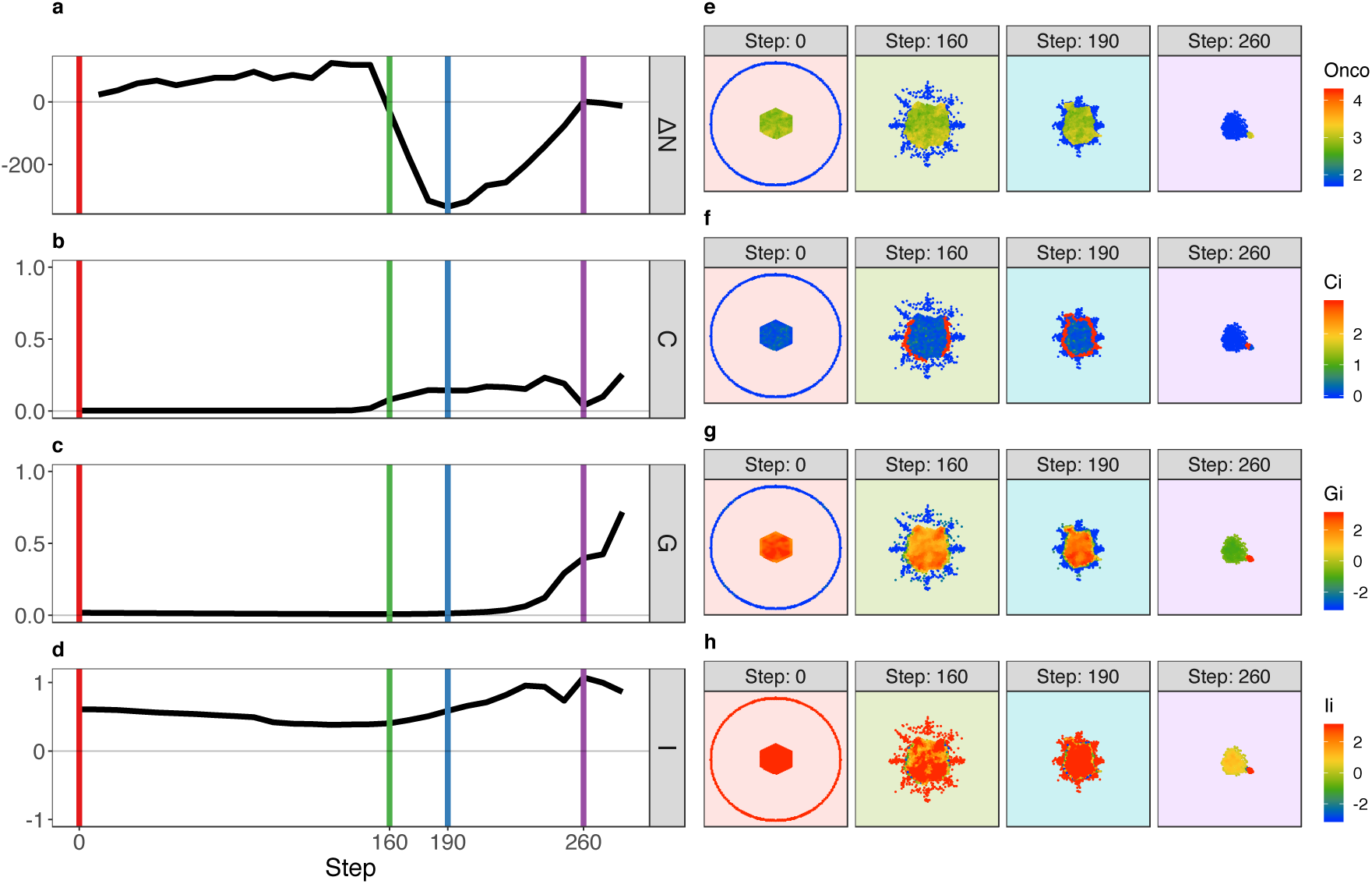
Predation model geostatistics. Values for the life of the model for a) the delta number of cancer cells from the step before; b) Geary’s C (Eq. 6); c) Getis-Ord G (Eq. 4); d) Moran’s I (Eq. 5). State of the model at time steps denoted by colored vertical lines in a-d for e) oncoprotein expression; f) Local Geary’s *C*_*i*_ (Eq. 9); g) Getis-Ord *G*_*i*_ (Eq. 7); h) Anselin Moran’s *I*_*i*_ (Eq. 8)

The local geostatistics in the predation model exhibit patterns dependent on area. During the initial predatory interaction in steps 160-190, we observe a marked increase in local spatial heterogeneity at the interaction interface (Figure 2f). As the space the cancer cells reside in is constrained, it becomes more of a spatial hotspot relative to the immune cell population surrounding it (Figure 2g). Overall, we can utilize local Geary’s *C*_*i*_ as a proxy for the presence of an ecological interaction and we observe a hotspot response that is dependent on area constraints.

### 4.2. Geostatistics stabilize to baseline after initial mutual interaction

The end stage global spatial behavior of the mutualism model looks very similar to the initialization state (Figure 3 b-d). Initially, there is relatively low spatial heterogeneity in the two tumors as evidenced by a low value for Geary’s *C* and a high value for Moran’s *I* (Figure 3b,d). As the model progress, ΔN steadily increases in line with the size of the tumors until step 130 when the two tumors begin a mutualistic relationship. Increased spatial heterogeneity is observed through increased local Geary’s *C*_*i*_ at the interface between the two tumors (Figure 3f) and is visualized globally by increases in Geary’s *C* and Moran’s *I*. The ΔN stalls during this initial interaction as the space available for each tumor to grow into has been decreased (Figure 3e-h). However, after an increase in proliferative capacity due to the mutualistic interaction, ΔN continues an upward trajectory. While we observe a similar increase in measured global spatial heterogeneity as the predation model, we observe very different global spatial clustering and local spatial patterns that differ greatly between the two models.

**Fig. 3.**
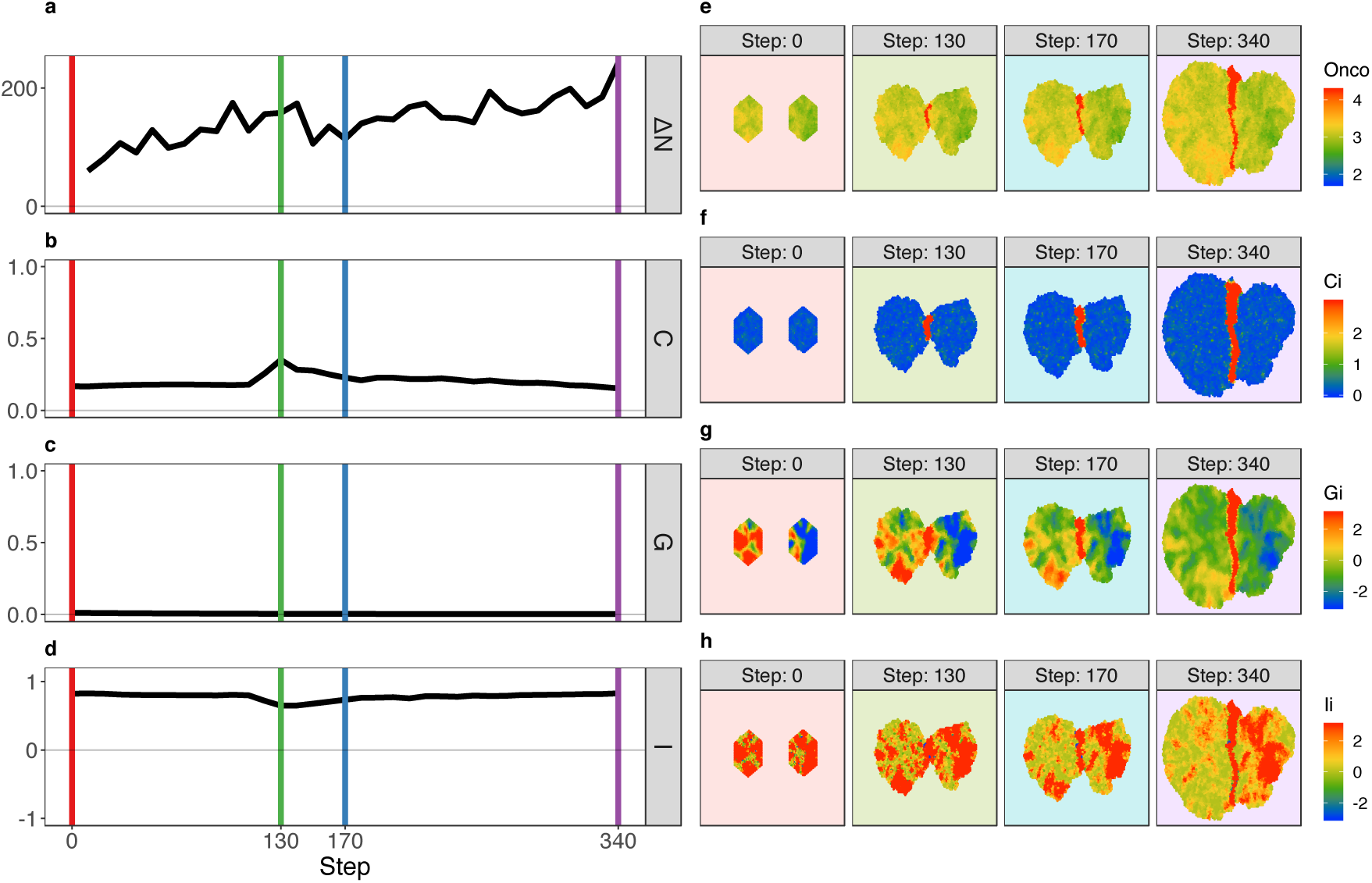
Mutualism model geostatistics. Values for the life of the model for a) the delta number of cancer cells from the step before; b) Geary’s C (Eq. 6); c) Getis-Ord G (Eq. 4); d) Moran’s I (Eq. 5). State of the model at time steps denoted by colored vertical lines in a-d for e) oncoprotein expression; f) Local Geary’s *C*_*i*_ (Eq. 9); g) Getis-Ord *G*_*i*_ (Eq. 7); h) Anselin Moran’s *I*_*i*_ (Eq. 8)

For the rest of the model’s life we observe a return to the baseline low heterogeneity global geostatistics even though there is a clear increase in local spatial heterogeneity. The increase in local heterogeneity is matched by the same increase in overall deviation of oncoprotein expression from the mean, resulting in global geostatistics that behave as if there is low heterogeneity (Eqs. 5 and 6). This model highlights the utility of visualizing the global and local geostatistics simultaneously to accurately capture the spatial phenomena occurring in the tumor.

### 4.3. Commensal interaction results in a new geostatistical baseline

The commensalism model displays similar global spatial behavior to the mutualism up through the initial interspecific interaction with low spatial heterogeneity that increases when the two tumors meet as evidenced by an increase in Geary’s *C* and decrease in Moran’s *I* (Figure 4b,d). However, as the model progresses, we do not observe the same return to baseline heterogeneity that the mutualism model displayed, but instead a new global pattern that more accurately represents the local spatial heterogeneity (Figure 4e-h) and is able to discriminate from the mutualistic interactions. This can be explained by the smaller region affected by an increase in oncoprotein expression in the commensal model (Figure 4e) relative to the mutual model (Figure 3e), resulting in a smaller increase to the mean oncoprotein in the model leading to an increase in Geary’s *C* (Eq. 6) and decrease in Moran’s *I* (Eq. 5). Geostatistics that have no dependence on the mean, like local Geary’s C (Eq. 9) exhibit spatial phenomena (Figure 4f) similar to the mutual model. Counterintuitively, in this scenario, a smaller local spatial effect results in a larger global spatial effect. Once again highlighting the utility of visualizing local and global phenomena simulatenously.

**Fig. 4.**
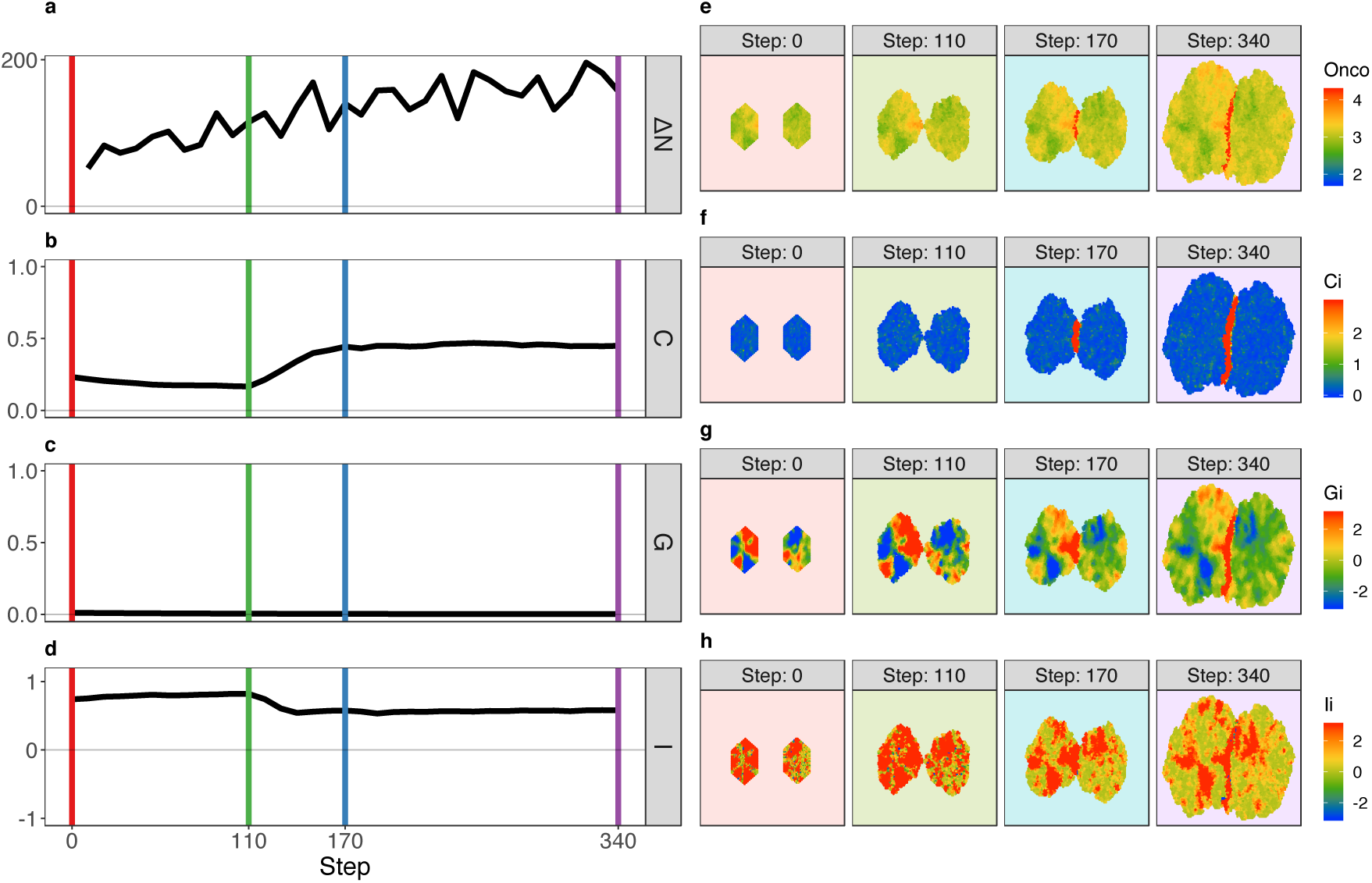
Commensalism model geostatistics. Values for the life of the model for a) the delta number of cancer cells from the step before; b) Geary’s C (Eq. 6); c) Getis-Ord G (Eq. 4); d) Moran’s I (Eq. 5). State of the model at time steps denoted by colored vertical lines in a-d for e) oncoprotein expression; f) Local Geary’s *C*_*i*_ (Eq. 9); g) Getis-Ord *G*_*i*_ (Eq. 7); h) Anselin Moran’s *I*_*i*_ (Eq. 8)

### 4.4. Geostatistics continually change after initial parasitic interaction

The parasitism model was initiated in the same manner as the mutualism and commensalism models and, as a result, we observe similar behavior through the initial interspecific parasitic interaction (Figure 5). However, we observe a much more drastic increase in global spatial heterogeneity (Figure 5b,d) in contrast to the mutual and commensal models. This behavior persists throughout the life of the model. The initial drastic change can be explained by a relatively small change in the mean oncoprotein expression due to an increase in one tumor matched by a decrease in the other, combined with a drastic increase local spatial interactions as we can observe clearly in the end stage spatial expression profiles at the tumor interface at step 340 in Figure 5e,h.

**Fig. 5.**
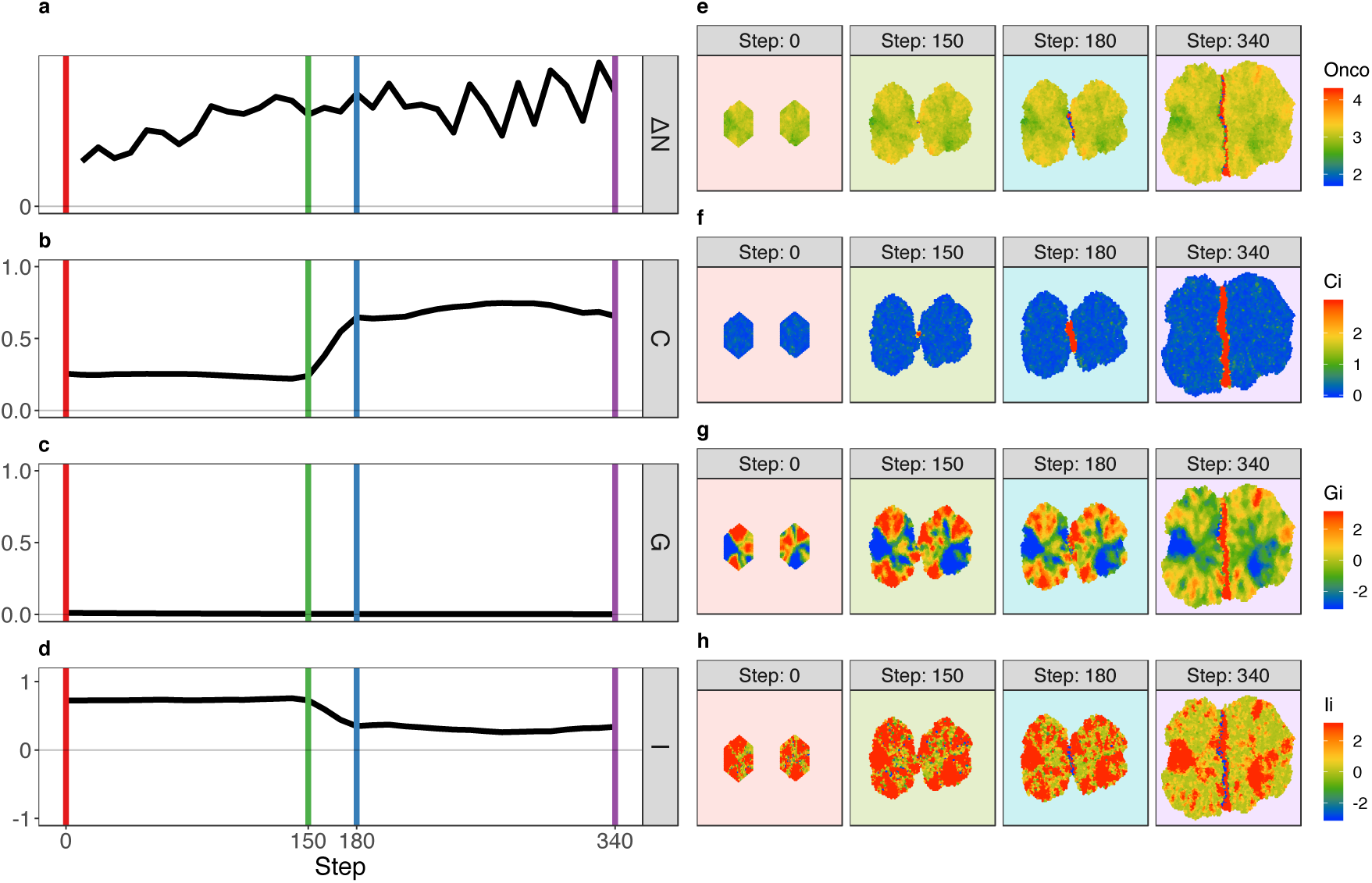
Parasitism model geostatistics. Values for the life of the model for a) the delta number of cancer cells from the step before; b) Geary’s C (Eq. 6); c) Getis-Ord G (Eq. 4); d) Moran’s I (Eq. 5). State of the model at time steps denoted by colored vertical lines in a-d for e) oncoprotein expression; f) Local Geary’s *C*_*i*_ (Eq. 9); g) Getis-Ord *G*_*i*_ (Eq. 7); h) Anselin Moran’s *I*_*i*_ (Eq. 8)

The parasitic relationship also results in greater variability of global phenomena over time with respect to ΔN and spatial heterogeneity (Figure 5a-d). This is because the change in fitness due to a change in oncoprotein expression is not symmetric about the mean(Eq. 3). More specifically, a positive change in oncoprotein expression that results in a doubling of replicative capacity has the effect of almost removing the ability to replicative when the change is negative. This results in global visual patterns that are very sensitive to the number of cells on either side of the parasitic relationship at a particular step. While this may be a peculiarity of how this model was written, it elucidates an interesting relationship between local cellular types and global patterns.

## 5. Limitations

Overall, we find that our proposed approach is able to effectively capture the heterogeneity and changes therein across a range of tumors exhibiting diverse ecological behaviors. However, we note that the methods have further room for improvement. One area that remains an ongoing challenge is with regards to edge effects. Relatively poor estimation of geostatistics at the outside borders of regions may arise from the same underlying stationary spatial process. However, as we are focusing on interactions between cells which, for the most part, occur in interior areas of the simulated tumors this is likely not a critical barrier in the short term. Along a similar vein, as the simulation progresses and the tumors become larger, the internal area grows exponentially faster than the border area; lessening the influence of border effects on determination of geostatistics. One additional challenge in visualizing tumor heterogeneity may arise when using a two-dimensional measurement to describe a subset of a three-dimensional process. Notably, the calculation of each geostatistic is agnostic to the dimensionality of space and the weighting matrix only captures which cells are neighbors and not their respective spatial distances. For profiling the ecological interactions of concern, a two-dimensional model is likely a sufficient representation. We additionally note that our tumor growth model is extremely simple. We don’t fully desribe vascularization or any of the other factors that might significantly alter tumor development and ecological interactions. Critically, our study is focused on the visualization and quantification aspects of tumor ecologies rather than on the details of tumor development. Consequently, our model is of sufficient complexity to test our approach to visualize tumor heterogeneity.

## 6. Conclusion

We have introduced a new method that uses a comprehensive geostatistical survey approach to examine the ecological interactions, heterogeneity of tumors and their local and global spatial patterns. The method was tested by growing virtual tumors using an agent-based model. Our approach is both sensitive to changing spatial patterns and stable with respect to stationary spatial processes. We also observe that quantifying and visualizing global and local spatial phenomena simultaneously is able to discriminate between different ecological relationships that single geostatistical measures are not. We see this model and visualization framework being used to test hypothesis of tumor evolution. For example, to characterize the spatial context differences between a successful and an ineffective immune response. Additionally, this framework can be readily extended to multiplexed analysis of many cancer markers simultaneously; allowing a deep analysis and visualization of spatial patterns.

